# A redox cycle with complex II promotes sulfide quinone oxidoreductase dependent H_2_S oxidation

**DOI:** 10.1101/2021.09.08.459449

**Authors:** Roshan Kumar, Aaron P. Landry, Arkajit Guha, Victor Vitvitsky, Ho Joon Lee, Keisuke Seike, Pavan Reddy, Costas A. Lyssiotis, Ruma Banerjee

## Abstract

The dueling roles of H_2_S as an endogenously synthesized respiratory substrate and as a toxin, raise questions as to how it is cleared when the electron transport chain is inhibited. Sulfide quinone oxidoreductase (SQOR) is a mitochondrial inner membrane flavoprotein that catalyzes the first step in the H_2_S oxidation pathway and uses coenzyme Q (CoQ) as an electron acceptor. However, complex IV poisoning by H_2_S inhibits complex III-dependent recycling of CoQH_2_, which is needed to sustain H_2_S oxidation. We have discovered that under these conditions, reversal of complex II activity using fumarate as an electron acceptor, establishes a new redox cycle with SQOR. The purine nucleotide cycle and the malate aspartate shuttle are sources of fumarate in H_2_S treated cells, which accumulate succinate. Complex II knockdown decreases the efficiency of H_2_S clearance and increases recovery time to the basal respiration rate in H_2_S treated cells. In contrast, attenuation of complex I, which is a major competitor for the mitochondrial CoQ pool, has the opposite effects. Targeted knockout of complex II in murine intestinal epithelial cells that are routinely exposed to microbiota derived H_2_S, decreases serum, urine, and fecal thiosulfate, a product of H_2_S oxidation. Our study identifies a metabolic reprogramming response to H_2_S that furnishes fumarate as an alternate electron acceptor and supports H_2_S oxidation independent of complex IV activity. Complex II-linked redox cycling of SQOR has important implications for gut H_2_S metabolism as colonocytes are routinely exposed to high concentrations of this gas derived from the microbiota.

**One Sentence Summary:** Reversal of complex II sustains and prioritizes H_2_S oxidation when respiration is poisoned.

---

The discovery of H_2_S as an endogenously synthesized signaling molecule in mammals has fueled a growing literature on its physiological effects (1). Mechanistic insights into how H_2_S modulates cellular responses are however, scarce (2, 3), and much attention has been focused on protein persulfidation, a reactive posttranslational modification of cysteine (4) that has been identified in hundreds of proteins (5, 6). On the other hand, the best characterized cellular effects of H_2_S are its oxidation via a dedicated mitochondrial pathway (7), and its inhibition of complex IV (8) in the electron transport chain (ETC), leading to respiratory poisoning (Fig. 1a). The sulfide oxidation pathway begins with the conversion of H_2_S to glutathione persulfide catalyzed by sulfide quinone oxidoreductase (SQOR), an inner mitochondrial membrane flavoprotein (9). Electrons released from H_2_S oxidation are transferred to coenzyme Q (CoQ) and enter the ETC at the level of complex III, making H_2_S an inorganic substrate for oxidative phosphorylation in mammals (10). The remainder of the pathway successively converts glutathione persulfide to thiosulfate and, in some cells, to sulfate (11). The role in signaling, if any, of the reactive sulfur species formed during H_2_S oxidation remains to be fully elucidated (12). In this study, we report that a noncanonical redox circuit is established when complex IV is inhibited, via reversal of complex II activity, which allows continued oxidation of H_2_S.

**Figure 1.**
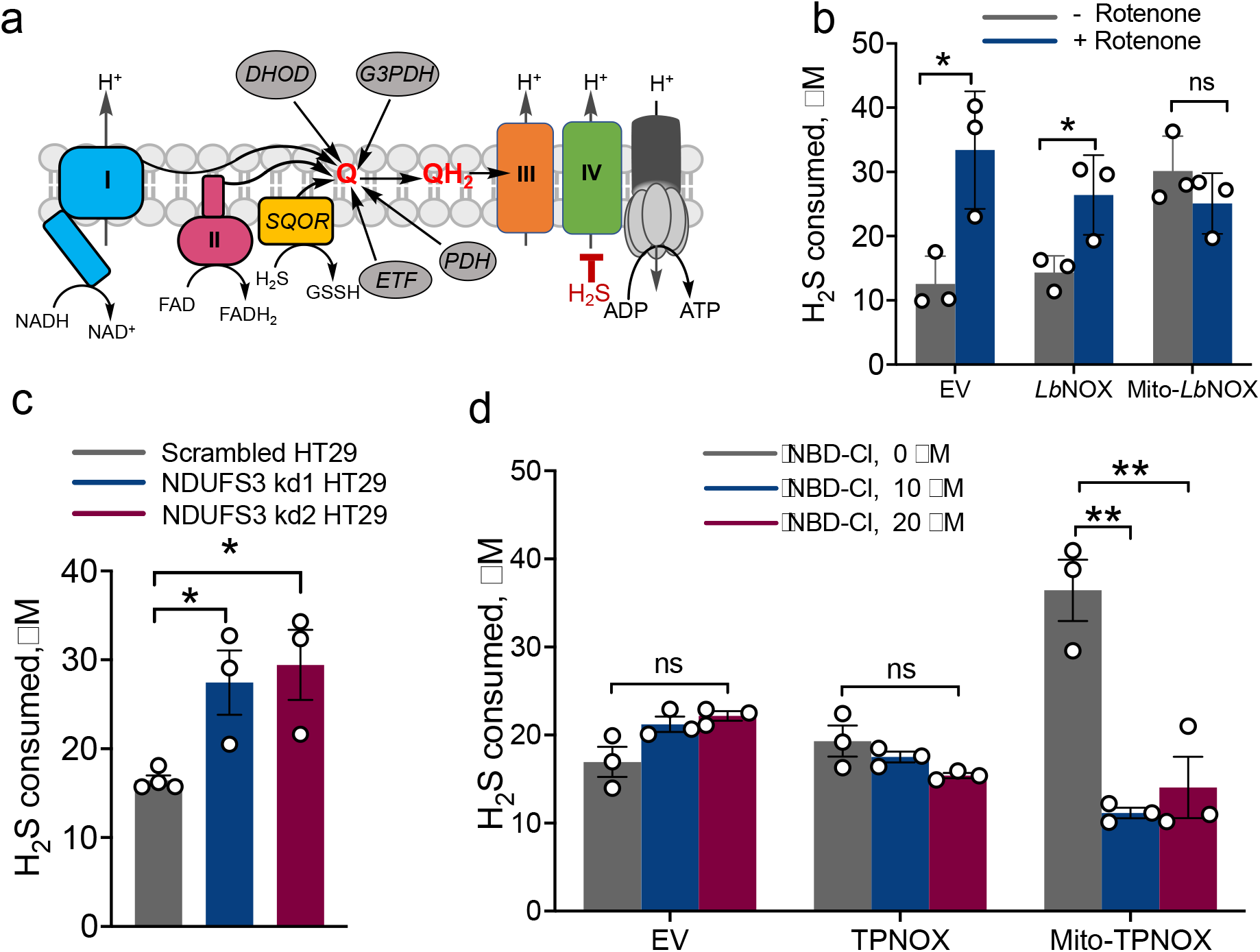
The mitochondrial NADH pool influences the efficiency of H_2_S oxidation in HT29 cells. (**a**) Scheme showing that multiple CoQ (Q) users compete with SQOR including complexes I and II, dihydroorotate dehydrogenase (DHOD), glycerol 3-phosphate dehydrogenase (G3PDH), proline dehydrogenase (PDH) and the electron transfer flavoprotein (ETF). (**b**) H_2_S oxidation is enhanced in cells expressing mitochondrial but not cytoplasmic *Lb*NOX versus the empty vector (EV) control. Rotenone (2 μM) enhanced H_2_S clearance in control and cytoplasmic but not mitochondrial expressing *Lb*NOX cells. (**c**) Disruption of complex I by NDUFS3 knockdown enhanced H_2_S oxidation. (**d**) Mitochondrial expression of TPNOX accelerates H_2_S oxidation, which is inhibited by NBD-Cl. The data are represent the SEM of 3 independent experiments (**p<0.001 and *p<0.05).

SQOR functions as a respiratory shield, sensitizing the ETC to H_2_S poisoning when its activity is attenuated (13). At low H_2_S concentrations however, SQOR activity increases respiration as measured by the oxygen consumption rate (OCR) (14). The dual potential to stimulate electron flux and inhibit the ETC, raises questions as to whether modulation of mitochondrial bioenergetics by H_2_S is pertinent to its cellular signaling mechanism and fans out to other compartments via redox and metabolomic changes (15).

SQOR is one of several consumers of CoQ (Fig. 1a) and sulfide oxidation is impaired in CoQ deficiency (16). SQOR activity has the potential to cause a reductive shift in the CoQ pool, particularly at H_2_S concentrations that partially or fully inhibit complex IV. H_2_S also indirectly perturbs the NAD^+^/NADH and FAD/FADH_2_ couples that are connected to CoQ/CoQH_2_ via the ETC. We have previously demonstrated that H_2_S induces a reductive shift in the NAD^+^/NADH redox couple, creating an electron acceptor insufficiency that leads to uridine and aspartate deficiency and enhances reductive carboxylation (13). While uridine limitation results from the CoQ dependence of dihydroorotate dehydrogenase in the pyrimidine pathway (Fig. 1a), aspartate deficiency results in part from reduced flux through the TCA cycle and the NADH-linked malate-aspartate shuttle. Furthermore, H_2_S stimulates the Warburg effect, enhancing glucose consumption and lactate production (17), and stimulates lipid biogenesis (18).

The effects of H_2_S on the ETC itself has received scant attention (10, 17, 19). The observed increase in succinate and decrease in malate at H_2_S concentrations that inhibit respiration was proposed to result from complex II reversal (10). While the same authors later proposed that H_2_S induces reverse electron transfer through complex I (14), neither model was evaluated experimentally. A recent study on oligomycin-treated murine microglia reported increased OCR upon exposure to an H_2_S donor and interpreted as evidence of reverse electron transfer through complex I (20). The known drivers of mitochondrial reverse electron transfer, which leads to reactive oxygen species (ROS) generation, are a high membrane potential and an over-reduced CoQ pool (21). Since respiratory poisons depolarize the membrane by limiting electron-coupled proton transfer (Fig. 1a), the premise for H_2_S-induced reverse electron transfer is unclear. Furthermore, the study contradicted the reported lack of H_2_S-induced ROS production (22).

Studies in our laboratory have focused primarily on colonic epithelial cells (13, 17, 18) that are routinely exposed to high concentrations of H_2_S from gut microbiota, estimated to range from ~0.2 to 2.4 mM (23, 24). In this study, we report that rewiring within the ETC circuitry via complex II reversal, sustains H_2_S oxidation under conditions of respiratory poisoning with fumarate serving as an electron acceptor. These results have important implications for understanding the mechanism by which intestinal epithelial cells respond to routine exposure to high H_2_S levels derived from the microbiota and potentially, the role of H_2_S in signaling a shift in energy metabolism.

## Results

### SQOR catalyzes sulfide-dependent reduction of O_2_

We examined whether O_2_ can serve an alternate electron acceptor for SQOR since complex IV poisoning by H_2_S should not restrict O_2_ availability (Supplementary Fig. 1a). We found that in the presence of sulfide and sulfite but in the absence of CoQ, nanodisc-embedded SQOR (*nd*SQOR) (25) catalyzed O_2_ consumption (Supplementary Fig. 1b). From the linear dependence of OCR on O_2_ concentration, a *k*_on_ of 3370 ± 290 M^−1^ s^−1^ was estimated (Supplementary Fig. 1c). Oxygen (*k* ~14 min^−1^ at 75 μM O_2_) is however, a significantly less efficient electron acceptor than CoQ (15 × 10^3^ min^−1^ at 75 μM CoQ) (26).

In the presence of a slight excess of sulfide (10 μM) and sulfite (15 μM), SQOR (7.5 μM) catalyzed the consumption of an equimolar concentration of O_2_ (7.3 ± 0.6 μM). This reaction stoichiometry predicted that the products of O_2_ reduction by SQOR could be either 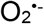 and FADHo or H_2_O_2_ and FAD. The equivalence between the concentration of H_2_O_2_ produced (7.6 ± 0.6 μM) and O_2_ consumed is consistent with the two-electron reduction of O_2_ by SQOR (Supplementary Fig. 1a). The concentration of H_2_O_2_ was significantly diminished (0.2 ± 0.1 μM) when catalase was added to the reaction mixture. The approximately 1:1:1 stoichiometry of sulfide added:O_2_ consumed:H_2_O_2_ produced is consistent with electron transfer from FADH_2_ to O_2_ via a C4a-hydroperoxy FAD intermediate (Supplementary Fig. 1d), as proposed in other O_2_-activating flavoenzymes (27).

### Complex I activity decreases the efficiency of H_2_S oxidation

Complex I-dependent oxidation of NADH with concomitant reduction of CoQ, is a major source of electron flux in the ETC and is expected to influence the efficiency of H_2_S oxidation. We have previously reported that H_2_S causes a reductive shift in the NAD^+^/NADH ratio by inhibiting complex IV (13). H_2_S oxidation was unaffected by the cytoplasmic, but significantly enhanced by the mitochondrial expression of the water forming NADH oxidase, *Lb*NOX (28) (Fig. 1b). Rotenone, a complex I inhibitor, increased H_2_S oxidation in control and *Lb*NOX but not mito*-Lb*NOX cells (Fig. 1b). Knockdown of NDUFS3 (Supplementary Fig. 2), which is required for complex I assembly, increased H_2_S oxidation (Fig. 1c). Collectively, these data demonstrate that the cellular H_2_S oxidation capacity can be limited by the mitochondrial NADH pool.

The mitochondrial NADH and NADPH pools are interconnected via the activity of the electrogenic nicotinamide nucleotide transhydrogenase (NNT) located in the inner mitochondrial membrane. Cytoplasmic expression of TPNOX, a genetically encoded water forming NADPH oxidase (29), had no effect on H_2_S oxidation, while mitochondrial expression enhanced clearance (Fig. 1d). The NNT inhibitor NBD-Cl (4-chloro-7-nitrobenzofurazan chloride) attenuated the mito-TPNOX effect, further demonstrating that the capacity for cellular H_2_S oxidation is linked to the status of the mitochondrial NAD(P)H redox pool (Fig. 1d).

### Succinate accumulates in response to H_2_S

Metabolomics analysis after exposure to Na_2_S (100 μM, 1 h) revealed a number of changes in glycolytic, TCA cycle (13) and purine metabolism intermediates in malignant HT29 cells (Fig. 2a,b). Interestingly, H_2_S treatment led to ~5.5-fold higher levels of succinate. To test whether succinate accumulation resulted from reversal of complex II activity (Fig. 2c), we used dimethyl fumarate (DMF), a membrane permeable derivative of fumarate that increases intracellular fumarate concentration (30). DMF accelerated H_2_S oxidation in 4 out of 5 colorectal carcinoma lines but not in RKO cells (Fig. 2d and Supplementary Fig. 3). Two other complex II inhibitors, dimethyl malonate and dimethyl itaconate, also inhibited H_2_S clearance, while diethyl succinate did not (Supplementary Fig. 4). Knocking down SDHA (Supplementary Fig. 5), the complex II subunit that catalyzes the reversible oxidation of succinate to fumarate, reduced H_2_S clearance (Fig. 2e). DMF shortened the recovery time for return to basal OCR following respiratory inhibition by H_2_S in HT29 (Fig. 2f-h), HCT116, LoVo and DLD cells (Supplementary Fig. 6) but had no effect when SDHA was knocked down in HT29 cells (Supplementary Fig. 7). Together, these data are consistent with the model that H_2_S oxidation is facilitated by reversal of complex II activity.

**Figure 2.**
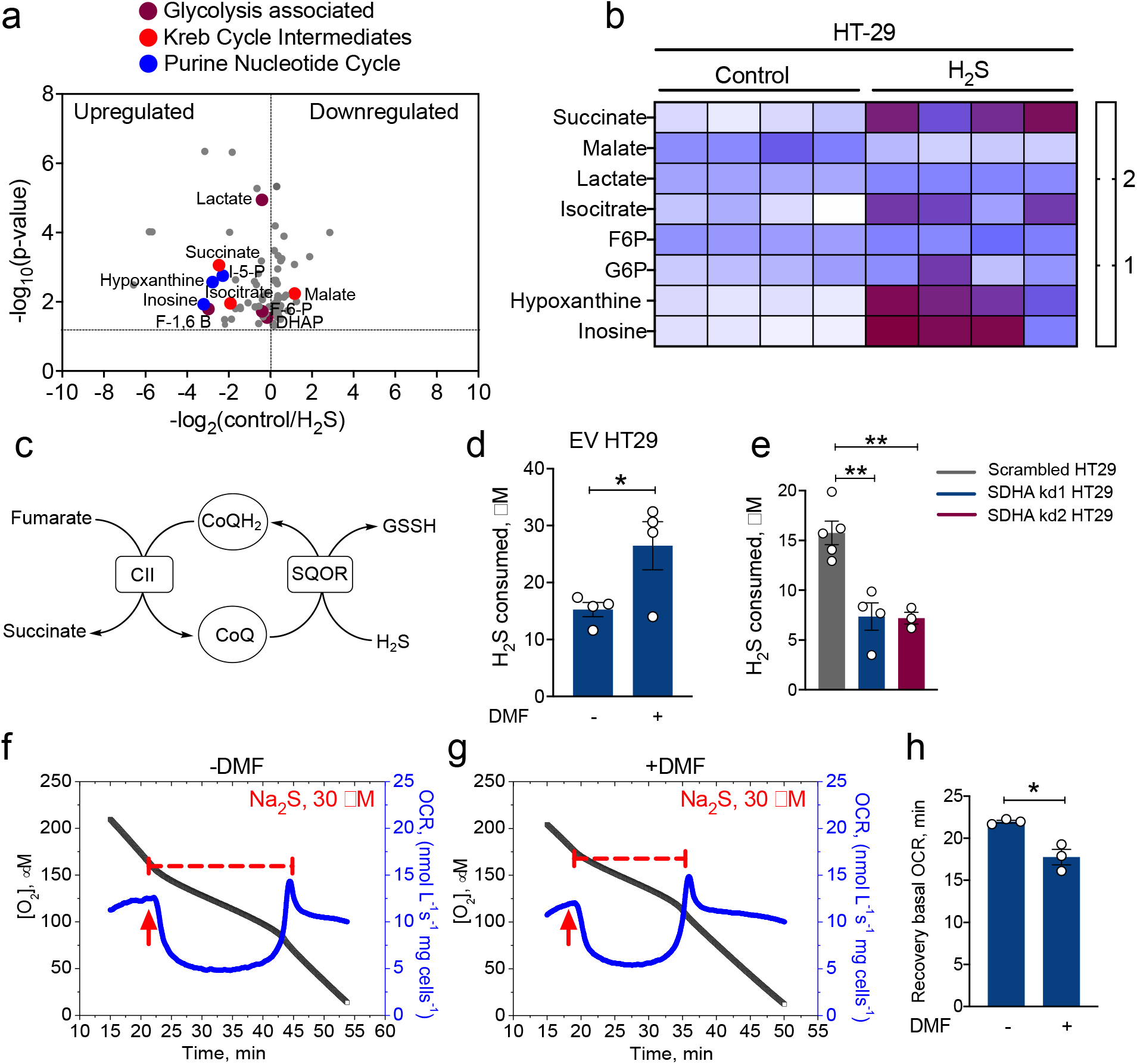
H_2_S induces succinate accumulation through reversal of complex II activity. (**a**) Volcano plot showing metabolite changes in response to Na_2_S (100 μM) treatment of HT29 cells for 1 h. (**b**) Heat map showing H_2_S-induced changes in select metabolites in HT29 cells. (**c**) Scheme showing how complex II reversal can regenerate CoQ for H_2_S oxidation. (**d**) DMF (100 μM) increases H_2_S oxidation in EV HT29 cells. (**e**) SDHA knockdown in HT29 cells reduces H_2_S oxidation. **(f,g)** The duration of respiratory inhibition in HT29 cells by H_2_S is longer in the absence (f) versus presence (g) of DMF (200 μM). The red arrows indicate when H_2_S (30 μM) was added. (**h**) Comparison of the time required by HT29 cells to return to the basal respiration rate ± DMF. The data in (d) and (e) represent the mean ± SEM of 3-4 independent experiments.

### The effect of complexes I and II on H_2_S-dependent OCR

To further test the influence of complexes I and II on the cellular response to H_2_S, OCR was monitored in control versus NDUFS3 and SDHA knockdown cells. NDUFS3 knockdown decreased basal OCR 2-fold (Fig. 3), consistent with complex I being a major entry point for electrons into the ETC. At a low concentration of H_2_S (10 μM), OCR activation in NDUFS3 knockdown cells was robust, and the peak increase in OCR was higher than in control and SDHA knockdown cells (Supplementary Fig. 8). At a higher H_2_S (20 μM) concentration, differences between the cell lines were clearly visible (Fig. 3a,b,c). While the NDUFS3 knockdown showed robust activation of OCR in response to H_2_S, the control and SDHA knockdown cells showed signs of inhibition. The SDHA knockdown cells took a longer time to recover basal OCR compared to controls. Following the first and second 20 μM H_2_S injection, control and SDHA knockdown cells showed signs of partial and severe respiratory inhibition, respectively, in contrast to NDUFS3 knockdown cells. At a higher H_2_S concentration (30 μM), control and SDHA knockdown cells responded with net inhibition of oxygen consumption in comparison to NDUFS3 knockdown cells, which exhibited a mixed response (Fig. 3d-f). These results indicate that the CoQ pool limits sulfide clearance and, in the absence of competition from complex I, cells clear sulfide more efficiently. The data also reveal that complex II has the opposite effect, i.e., it is advantageous for sulfide clearance, consistent with our model that complex II reversal supports H_2_S oxidation by catalyzing CoQH_2_ oxidation.

**Figure 3.**
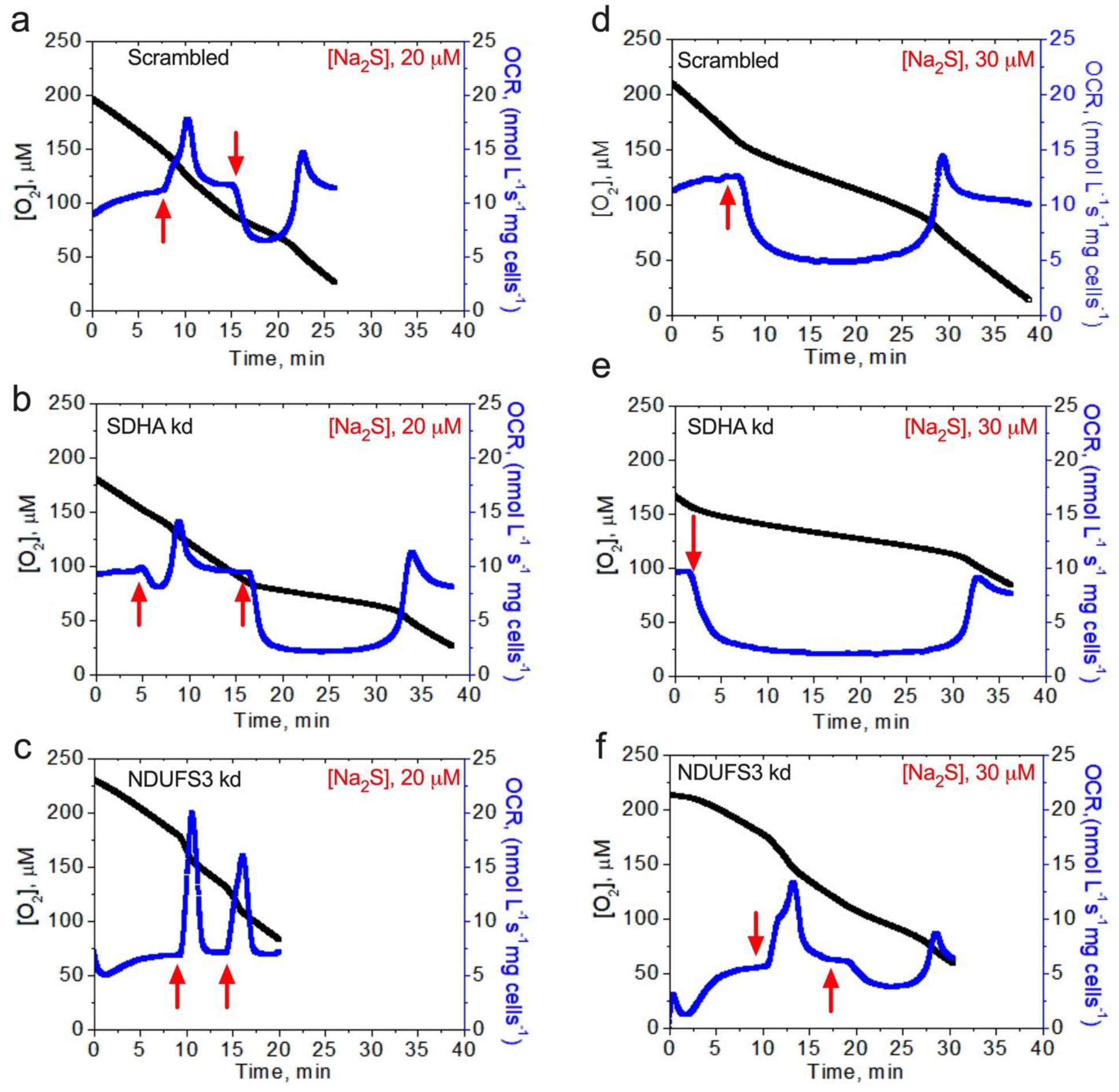
Complexes I and II influence H_2_S-linked OCR. Comparison of OCR activation with H_2_S (20 or 30 μM) in **(a, d)** scrambled, **(b, e)** SDHA knockdown, and **(c, f)** NDUFS3 knockdown HT29 cells. Red arrows indicate when H_2_S was added. The traces are representative of 3-5 independent experiments.

### Malate-aspartate shuttle and PNC furnish fumarate in H_2_S treated cells

Since the malate-aspartate shuttle and the purine nucleotide cycle (PNC) (Fig. 4a,b) are metabolic sources of fumarate in ischemic cells (21), we tested whether they also contribute to fumarate when the ETC is inhibited by H_2_S. For this, GOT1 and GOT2 (glutamic-oxaloacetic aminotransferases 1 and 2) expressed in the cytoplasm and mitochondrion, respectively, were knocked down in HT29 cells (Supplementary Fig. 9). GOT1 but not GOT2 knockdown increased H_2_S oxidation by ~38% compared to control cells (Fig. 4c). GOT1 knockdown also promoted H_2_S clearance as reflected by the shorter recovery time to the basal respiration rate (Supplementary Fig. 10). Inhibition of adenylosuccinate lyase with AICAR (5-aminoimidazole-4-carboxamide ribonucleotide) decreased H_2_S clearance by ~50% (Fig. 4d), consistent with a role for the PNC in this process.

**Figure 4.**
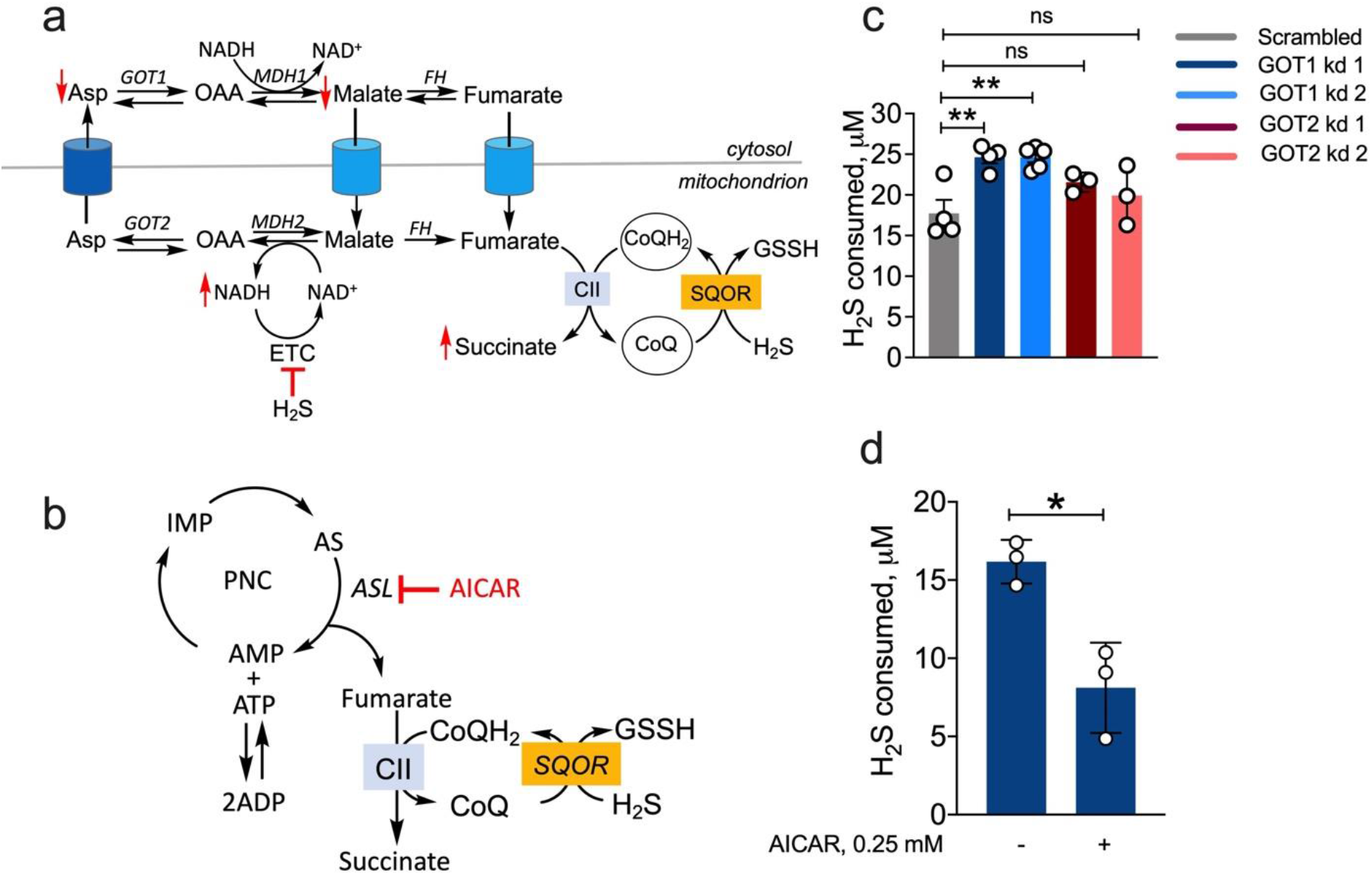
The PNC and the malate-aspartate shuttle support fumarate driven H_2_S oxidation. (**a,b**) Schemes showing that the malate-aspartate shuttle (a) and the PNC (b) are fumarate sources and that AICAR inhibits adenylosuccinate lyase (ASL). MDH1/2, FH, OAA and CII denote malate dehydrogenase 1/2, oxaloacetate, fumarate hydratase and complex II, respectively. (**c**) H_2_S oxidation is stimulated in GOT1 knockdown but unaffected by GOT2 knockdown in HT29 cells. (**d**) AICAR (0.25 mM) inhibits H_2_S clearance. The data in (c) and (d) represent the SEM of 3-4 independent experiments (*p<0.05).

### SDHA knockout in murine intestinal epithelial cells decreases H_2_S oxidation

To assess the physiological relevance of our observation that H_2_S clearance is supported by complex II working in reverse, we measured the impact of attenuating complex II on organismal H_2_S metabolism. For this, mice harboring loxP-flanked *Sdha* were crossed to mice expressing Cre recombinase under control of the villin promoter to specifically target intestinal epithelial cells, to generate *Vil1-Cre Sdha*^fl/fl^ *(Sdha^ΔIEC^)* mice as described previously (31). The rationale for targeting intestinal epithelial cells is that they are routinely exposed to high concentrations of H_2_S (23, 24) and actively oxidize sulfide (13). Thiosulfate, a stable product of H_2_S oxidation (Fig. 5a), is a stable biomarker of H_2_S metabolism. H_2_S on the other hand, is difficult to measure due to its volatility and low steady-state concentrations in biological samples (32). *Sdha^ΔIEC^* mice showed lower thiosulfate levels compared to control *Sdha*^fl/fl^ (Fig. 5b-d) revealing that the loss of complex II in intestinal cells caused local (feces) and systemic (serum and urine) perturbations in H_2_S oxidation.

**Figure 5.**
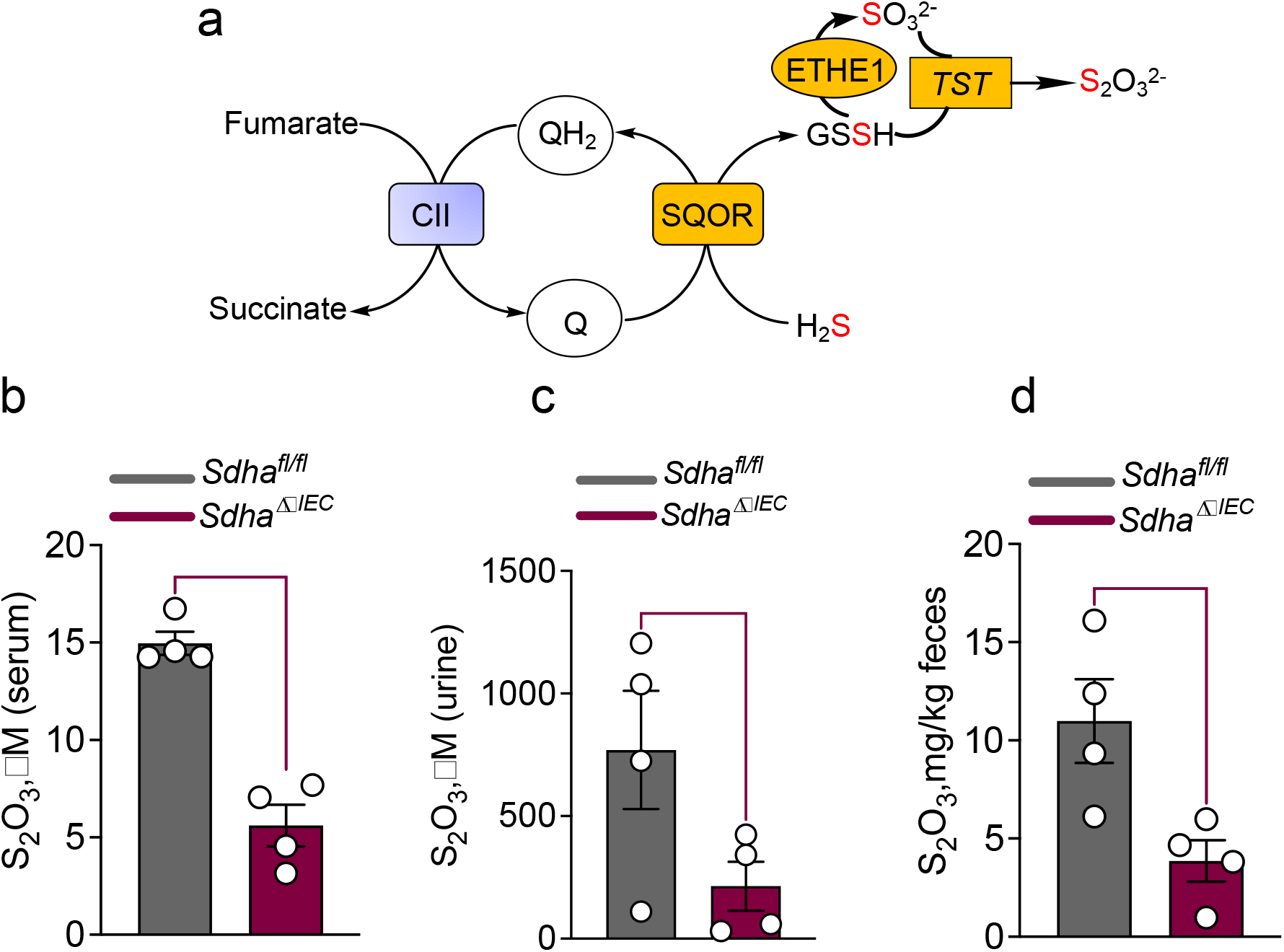
VillinCreSDHAfl/fl mice have reduced thiosulfate levels. (**a**) Scheme connecting H_2_S oxidation to thiosulfate production. (**b,c,d**) Quantitation of thiosulfate levels in control and *Villin^Cre^Sdha* knockout versus control (*Sdha*^fl/fl)^ mice in serum (b), urine (c) and feces (d). The data represent the SEM for samples collected from 4 mice in each group (*p<0.05).

## Discussion

In this study, we have uncovered a new mechanism for clearing H_2_S when its concentrations rise to levels that inhibit complex IV and preclude the use of O_2_ as the terminal electron acceptor for SQOR-dependent H_2_S oxidation. Such conditions might be relevant in the gut epithelium (where H_2_S exposure is high) or in ischemia (where O_2_ supply is cut off). Reversal of complex II activity under such conditions supports SQOR-dependent H_2_S oxidation, using fumarate as an alternate electron acceptor.

Metabolomic changes in HT29 cells in response to H_2_S provided clues to reprogramming driven changes that could potentially impact its clearance. Hypoxanthine and succinate, classic ischemic biomarkers (21, 33), also accumulate in response to H_2_S (Fig. 2b). Ischemic succinate accumulation is derived from oxidative TCA cycle metabolism (34) as well as from complex II-catalyzed reduction of fumarate (21). Fumarate is derived via the malate-aspartate shuttle and the PNC (21). Since H_2_S decreases the NAD^+^/NADH ratio and stimulates reductive carboxylation of α-ketoglutarate (13), the effect of the oxidative TCA cycle on H_2_S clearance was not examined. The PNC and the malate aspartate shuttle both impacted H_2_S clearance (Fig. 4c,d). The PNC is activated in response to a drop in the adenylate energy charge (35), and is consistent with lower ATP levels in H_2_S-treated cells (17) as well as the observed increase in inosine, which is formed via deamination of adenosine.

Knockdown of GOT1, but not GOT2, increased the efficiency of H_2_S clearance, suggesting that the cytoplasmic arm of the malate-aspartate shuttle is an important source of fumarate. H_2_S leads to aspartate deficiency (13), potentially stimulating GOT1-catalyzed transamination of oxaloacetate to aspartate rather than the reverse, which is consistent with lower malate levels in H_2_S-treated cells (Fig. 2b). In GOT1 knockdown cells, oxaloacetate should be more available for malate dehydrogenase catalyzed reduction to malate, which can be dehydrated to fumarate (Fig. 4a) by fumarate hydratase that is present in the cytoplasm and the mitochondrion (36). Cytosolic fumarate can potentially enter the mitochondrion via a dicarboxylate carrier (37).

Our studies support a model for efficient H_2_S clearance by SQOR when the H_2_S concentration is low with complexes I and II competing for the CoQ pool and complex III recycling CoQH_2_ (Fig. 6a). However, when H_2_S concentrations rise and inhibit complex IV, utilization of fumarate as an electron acceptor by complex II sustains recycling of CoQH_2_ (Fig. 6b). Complex II catalyzes the reversible oxidation of succinate to fumarate (38) and exhibits similar *K*_M_ values for both substrates (39, 40). Under *in vitro* assay conditions, the ratio of succinate oxidation to fumarate reduction catalyzed by the succinate dehydrogenase component of complex II varies substantially with the electron acceptor, and ranges from ~0.1 to 50 for succinate:fumarate consumed (40). Under physiological conditions, flux through the forward versus reverse reaction is governed by the concentration of the respective substrates and by the potentials of the relevant redox couples. In the mitochondrial matrix (pH ~7.7), the standard redox potential for the fumarate/succinate couple (E°’=+ 30 mV) is similar to that ubiquinone (+40-60 mV at pH 7.0, decreasing 60 mV per increase in pH unit (41)), but higher than of the FAD/FADH_2_ couple (−79 mV (42, 43)). The reversibility of complex II in cells is supported by its ability to sustain proficient growth on fumarate as a terminal electron acceptor when expressed under anaerobic conditions in an *E. coli* strain lacking fumarate reductase (44). These data support the plausibility of complex II reversal under conditions when the ETC is blocked, and the CoQ pool is over-reduced.

**Figure 6.**
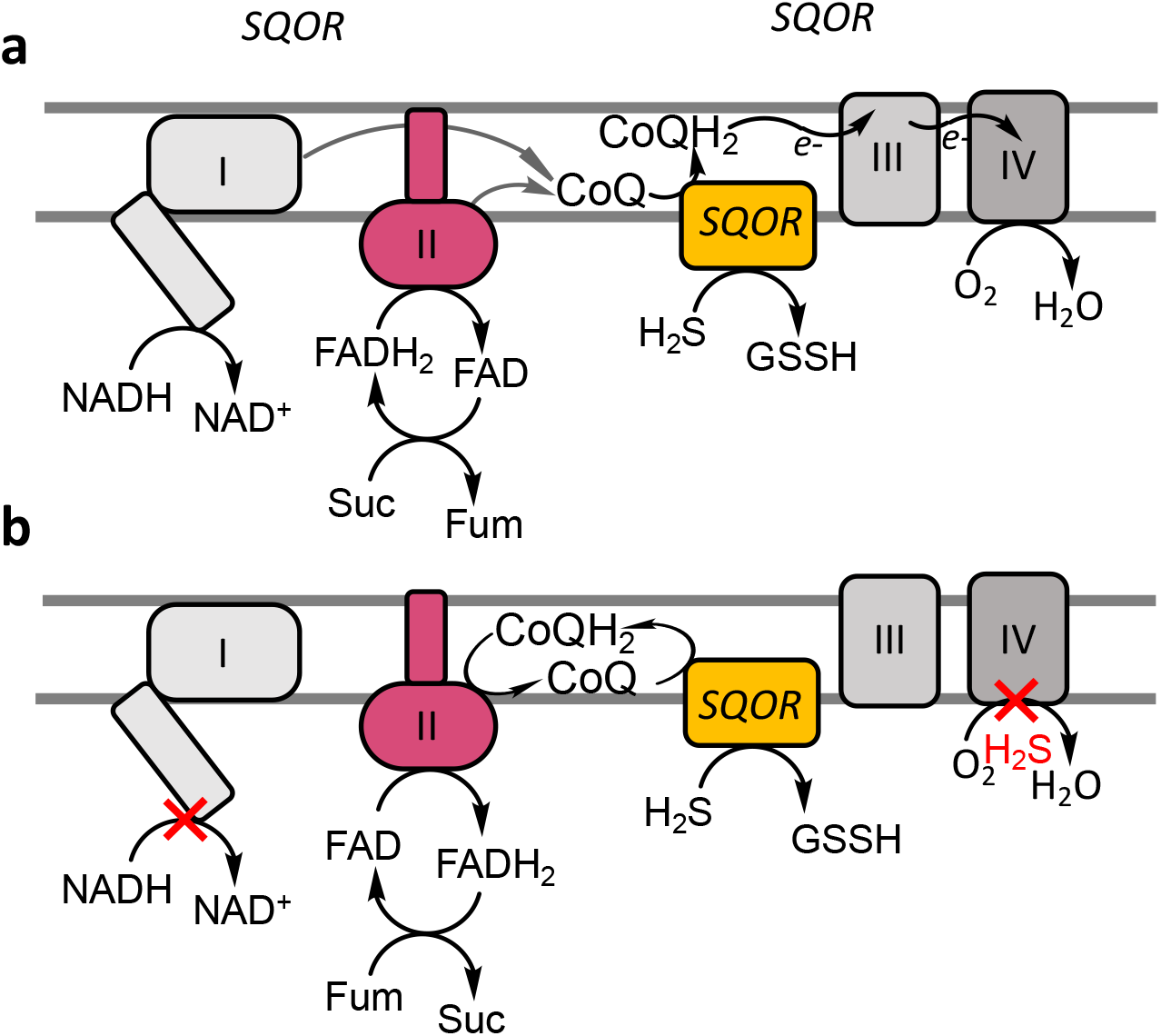
Alternate redox cycles for disposing H_2_S. (**a**,**b**) CoQH_2_ formed during H_2_S oxidation and by complexes I and II, enters the ETC at the level of complex III (a). When complex IV is inhibited by H_2_S, blocking recycling of CoQH_2_ by complex III, CoQH_2_ can be oxidized by complex II, concomitant with fumarate reduction and succinate accumulation (b).

Modulation of H_2_S metabolism by complex I was demonstrated by its inhibition by rotenone and by NDUFS3 knockdown; both enhanced H_2_S clearance (Fig. 1b,c), as expected, and is consistent with their increased sulfide-induced OCR compared to control cells (Fig. 3). On the other hand, complex II inhibition (with DMM or DMI) or SDHA knockdown, decreased the efficiency of H_2_S clearance while DMF increased it (Fig. 2, Supplementary Figs. 3,4). Under conditions of complete coupling, for every mole of sulfide oxidized by SQOR, ETHE1 and complex IV are predicted to consume 1 and 0.5 mole of O_2_, respectively. ETHE1 is a mononuclear iron-dependent persulfide dioxygenase, which catalyzes the conversion of glutathione persulfide to sulfite (45, 46). SDHA knockdown cells exhibited increased sensitivity to H_2_S-induced inhibition of OCR and took longer to recover, while DMF reduced the time to recovery of the basal OCR (Figs. 2 and 3). Collectively, these results support our model of complex II-dependent recycling of CoQH_2_ (Fig. 6b). It is important to note however, that interfering with complex II reduces but does not completely block H_2_S consumption. Thus, other mechanisms including SQOR-dependent reduction of O_2_ (Supplementary Fig. 1) might contribute to H_2_S removal.

The significant decrease in thiosulfate upon silencing SDHA in murine intestinal epithelial cells (Fig. 5) is notable for three reasons. It supports the physiological relevance of reverse complex II activity for H_2_S oxidation as loss of the canonical succinate oxidation activity would be expected to stimulate SQOR-dependent H_2_S oxidation by decreasing competition for the CoQ pool. Second, the observed change in thiosulfate levels in *Sdha*^*ΔIEC*^ mice reflect the impact of complex II on endogenous sulfide metabolism. Third, changes in urine and serum thiosulfate in *Sdha*^*ΔIEC*^ mice reveal the systemic impact of altered H_2_S metabolism at the host-microbe interface, which warrants further study.

We speculate that H_2_S-fueled succinate accumulation could have downstream metabolic effects. Succinate is a competitive inhibitor of α-ketoglutarate-dependent dioxygenases and its accumulation could broadly impact histone and DNA methylations (47). Furthermore, succinylation, a posttranslational modification of proteins (48), could be enhanced by H_2_S-driven succinate accumulation. Over >750 protein targets of succinylation have been identified, which are concentrated in mitochondria but also present in other compartments (49) and reversed by the NAD^+^−dependent sirtuin, Sirt5 (50). Succinylation reportedly increases complex II activity (49). We speculate that succinylation could be enhanced by the opposing effects of H_2_S on the succinate and NAD^+^ pools, in an autocorrective loop for activating complex II and prioritizing its removal.

In summary, our study reveals that metabolic reprogramming leads to the establishment of a new redox cycle between SQOR and complex II, permitting sustained H_2_S clearance. In addition to its relevance at the gut host-microbe interface, this circuitry could be important in the context of ischemia reperfusion injury. H_2_S is cytoprotective when administered at the time of reperfusion, reducing infarct size, inhibiting myocardial inflammation and preserving mitochondrial integrity (51). The rapid reoxidation of succinate that accumulates in the ischemic phase, drives ROS production during reperfusion (21). We posit that the cytoprotective effects of H_2_S could derive from its twin effects on complex IV inhibition and complex II reversal, thereby attenuating succinate-dependent ROS generation during reperfusion. Another cellular context in which H_2_S mediated ETC rewiring might be relevant is during the transition from a quiescent to proliferative state. While quiescent cells primarily rely on the high energy yield of oxidative phosphorylation, proliferating cells increase aerobic glycolysis to meet their energy needs and redirect mitochondrial metabolism for macromolecular precursor synthesis (52). The potential for H_2_S to function as an endogenous modulator of energy metabolism could be significant in this context and needs to be further understood.

## Acknowledgements

This work was supported in part by the grants from the National Institutes of Health (GM130183 to RB, NCI R01CA244931 to CAL and HL152605; HL149633; CA203542 to PR) and the American Heart Association (826245 to RK).

## Author contributions

RK-kinetics of H_2_S consumption, generation and analysis of TPNOX, NDUFS3 and SDHA knockdown cell lines thiosulfate in *Sdha*^Δ/IEC^ and control mice; APL-analysis of SQOR reactions; AG-OCR data generation and analysis; VV, HJL, CAL-metabolomics data generation and analysis; SK and PR-generation of *Sdha*^Δ/IEC^ mice; RK, APL and RB drafted the manuscript and all authors edited and approved the final version of the manuscript.

## Supplementary Materials

Materials and Methods

Fig S1 – S10

Supplementary References

## Data and materials availability

All data are available in the main text or the supplementary materials.

