## Supplementary for "A redox cycle with complex II promotes sulfide quinone oxidoreductase dependent H_2_S oxidation"

**This PDF file includes:**

Materials and Methods
Figs. S1 to S10

### Materials and Methods

***Materials*-**Sodium sulfide nonahydrate (431648), sodium sulfite (S0505), sodium selenite (S5261), CoQ (C7956), dimethyl malonate (63380), dimethyl itaconate (592498), diethyl succinate (8.00680), rotenone (R8775), dimethyl fumarate (242926), 4-chloro-7-nitrobenzofuran (163260), doxycycline (D3447), puromycin (P8833), protease inhibitor cocktail for mammalian tissue extract (P8340), RIPA lysis buffer (R0278) and apo-transferrin (T1147) were from Sigma. RPMI 1640 (11875-093), DMEM (11995-065), FBS (10437-028), trypsin-EDTA (25300-054), penicillin-streptomycin (15140-122), geneticin (10131-035), M199 (11150-059), epidermal growth factor (PHG0311), PBS (10010-023), DPBS (14040-133), and insulin (12585014) were from Gibco. Anti-Flag (20543-1-AP), anti-NDUFS3 (15066-1AP), anti-SDHA (14865-1AP), anti-GOT1 (14886-1AP), and anti-GOT2 (14800-1AP) antibodies were from Proteintech and the secondary anti-rabbit horseradish peroxidase-linked IgG antibody (NA944V) was from GE Healthcare.

***Assays for ndSQOR-catalyzed $O_2$ consumption and $H_2O_2$ production***

Human SQOR was purified and embedded in nanodiscs as described previously (1). $O_2$ consumption by $FADH_2$ in *nd*SQOR was monitored using an O2k respirometer (Oroboros Instruments, Austria), equipped with two polarographic $O_2$-sensing electrodes housed in separate 2 ml chambers. Each chamber was filled with 100 mM potassium phosphate, pH 7.4, and sulfide (100 μM) and sulfite (200 μM) were added before sealing the chambers and pre-incubating for ~5 min at 25 °C. The reaction was initiated by injecting *nd*SQOR (100 nM) and monitored over a period of ~10 min. Initial $O_2$ concentrations were varied by aerating $N_2$-purged buffer in the chambers before sealing when the desired $O_2$ concentration was reached. $H_2O_2$ production was assayed using the Pierce Quantitative Peroxide Assay Kit (ThermoFisher) according to the manufacturer's protocol.

***Cell culture***

HT29 cells were maintained in RPMI 1640 medium. HCT116, LoVo, DLD and RKO were maintained in DMEM medium. Both RPMI and DMEM media were supplemented with 10 % FBS, 100 units/ml penicillin and 100 µg/ml streptomycin. HCEC cells were cultured as described previously (2). All cells were maintained at 37 $^{o}$C with ambient $O_2$ and 5% $CO_2$ except HCEC, which were maintained at 2% $O_2$ and 5% $CO_2$.

***Ectopic expression of LbNOX and TPNOX***

*Lb*NOX and mito-*Lb*NOX and pINDUCER empty vector were obtained from Addgene. The pLVX-TRE3G empty vector, TPNOX, mito-TPNOX, and pLVX TET ON were a generous gift from Dr. Valentin Cracan (Scintillon Institute). The construction of HT29 cell lines stably expressing *Lb*NOX, mito-*Lb*NOX, TPNOX and mito-TPNOX have been described previously (3,4). Before the start of an experiment, these cells were incubated for 24 h with 300 ng/ml doxycycline to induce *Lb*NOX expression. The cells were routinely cultured in RPMI 1640 medium supplemented with 10% FBS, 100 units/ml penicillin, 100 µg/ml streptomycin and 300 µg/ml geneticin, and 1 µg/ml puromycin.

***Generation of shRNA mediated knockdown cells***

NDUFS3 and SDHA were targeted for knockdown using shRNA purchased from MISSION shRNA Library, Sigma. The clone IDs for NDUFS3 were: NM_004551.1-320s21c1 and NM_004551.1-628s21c1. The clone IDs for SDHA were: NM_004168.1-619s1c1 and NM_004168.1-1643s1c1. The doxycycline inducible GOT1 and GOT2 lentiviral constructs were from sub-cloned into the iDox-pLKO vector as described previously (5,6). Plasmids containing shRNA against specific genes or a scrambled sequence were submitted to the Vector Core (University of Michigan) for lentiviral packaging. For lentiviral infection, 7.5 x$10^{4}$ HT29 cells were seeded in a 6 well plate containing 2 ml per well of RPMI 1640 medium supplemented with 10

% FBS, 100 units/ml penicillin, and 100 µg/ml streptomycin. The transduction and selection protocols were the same as described for *Lb*NOX (3), and cells were selected with 1 µg/ml puromycin.

***Western blotting***

TPNOX expression in HT29 cells was monitored by growing cells in a 6 well plate for 24 h in RPMI 1640 medium as described above followed by a 24 h incubation with 300 ng/ml doxycycline. Then, the cells were washed with PBS twice before addition of 250 µl of RIPA lysis buffer containing 10 µl/ml protease inhibitor cocktail for mammalian tissue extracts and collected by scraping. Cells were frozen and thawed three times and centrifuged at 12,000 x *g* for 5 min. The protein concentration in the supernatant was measured using Bradford reagent (Bio-Rad). Protein lysates were similarly prepared from cells in which NDUFS3, SDHA and GOT1/2 were knocked down. Following separation by 10% SDS PAGE, proteins were transferred to a PVDF membrane and incubated overnight at 4 °C with primary anti-Flag antibody at a dilution of 1:1000 for TPNOX. Antibodies against NDUFS3, anti-SDHA, GOT1, and GOT2 (14800-1AP) were used at a dilution of 1:2000. Horseradish peroxidase linked anti-rabbit IgG was used at a dilution of 1:10000. Membranes were developed and visualized using the KwikQuant Digital-ECL substrate and imaging system.

***Cellular $H_2S$ consumption assay***

Cells were grown to ~90% confluency in 10 cm plates and on the day of experiment, washed with PBS and treated with 0.05% trypsin-EDTA (for ~10 min at 37 °C). Then, cells were resuspended in 10 ml complete media and centrifuged for 5 min at 4 $^{o}$C, 1700 x *g*. The cell pellet was resuspended in 1 ml modified DPBS (supplemented with 20 mM HEPES, pH 7.4, and 5 mM glucose) in a pre-weighed Eppendorf tube and centrifuged for 5 min at 4 $^{o}$C, 1700 x *g*. The supernatant was discarded, and the pellet weight was determined. Cells were suspended in modified DPBS to make a 5% cell suspension (w/v) in a 1 ml Eppendorf tube. When the effects of dimethyl malonate (DMM,10 mM) or dimethyl itaconate (DMI, 0.25 mM) were tested, cells were preincubated for 3 h with each reagent before making a 5% cell suspension in which the same concentration of DMM and DMI were included followed by addition of 100 µM $Na_2S$. Alternatively, when dimethyl fumarate (DMF, 100 µM) and diethyl succinate (DES, 5 mM) were tested, these reagents were added to a 5 % cell suspension in modified DPBS for 5 min prior to addition of 100 µM $Na_2S$. The suspension cultures were incubated at 37 °C with shaking (75 rpm). Samples (45 µl) were collected at time 0 and 10 min, mixed with 1 M Tris base (2.5 µl), and stored in dry ice. Control samples containing 10 mM DMM, 0.25 mM DMI, 100 µM DMF, or 5 mM DES and 100 µM $Na_2S$ in modified DPBS were incubated in parallel and the concentration of $H_2S$ lost from these sample was subtracted from the values obtained from the cell suspension samples containing the same reagents.

***Monobromobimane derivatization of sulfide and HPLC analysis***

The samples from the $H_2S$ consumption assay described above were thawed and mixed with 2.5 µl of 60 mM monobromobimane (in DMSO) and incubated in the dark at room temperature for 10 min followed by addition of 100 µl of metaphosphoric acid solution (16.8 mg/ml). The samples were vortexed and centrifuged for 5 min at 4 °C and 10,000 x *g*. The supernatant was collected in the dark and stored at -20 °C until further use. The samples were analyzed using a Zorbax Eclipse XDB-C18 column (5 µm, 4.6 x 150 mm, Agilent, CA) as described previously (7). Peaks were detected using excitation at 390 nm and fluorescence emission at 490 nm. A calibration curve with known concentrations of sodium sulfide was used to determine the concentration of $H_2S$ in samples.

***Metabolomics analysis***

Metabolomics analysis on HT29 cells treated ± 100 µM $Na_2S$ for 1 h was performed as described previously (2).

***OCR measurements***

Oxygen consumption was measured using the O2k respirometer. Cells were grown to ~90% confluency in 10 cm plates and on the day of experiment, washed with PBS, and then trypsinized with 1.5 ml of 0.05% trypsin-EDTA for ~10 min at 37 °C. Then, the cells were resuspended in 10 ml of complete medium and centrifuged for 5 min at 1700 x *g*, 4 °C. The cell pellet was resuspended in 1 ml of modified DPBS in a pre-weighed Eppendorf tube, the suspension was centrifuged for 5 min at 1700 x *g* and the weight of the pellet was recorded. The cells were suspended in modified DPBS to make a 5% cell suspension (w/v), which was stored on ice. At the start of the experiment, the cell suspension was diluted to 1% or 1.5% (for NDUFS3 knockdowns which showed lower basal OCR). The cell suspension was placed in the respirometer chamber and the OCR was allowed to stabilize over ~15-20 min at 37 °C with constant stirring at 750 rpm. $Na_2S$ (from a freshly prepared 10 mM stock solution in water) was injected into the sample to give the desired final concentration (10-30 µM) per injection.

***Mice***

*B6.Cg-Tg(Vil-cre)1000Gum/J* mice were purchased from the Jackson Laboratory (Bar Harbor, ME, USA). C57BL/6N-*Sdha*$^{tm2a(KOMP)Wtsi}$ mice were obtained from the Knock Out Mouse Project (KOMP) repository, University of California, Davis (Davis, CA, USA) and bred to *ACTFLPe* mice to excise the *FRT*-flanked region. The resulting *Sdha*$^{fl/fl}$ mice were bred to *Vil1-Cre* mice to create *Vil1-Cre Sdha*$^{fl/fl}$ (*Sdha*$^{\Delta/IEC}$) mice (8). Then,12-15 week-old mice were used in our experiments. The mice were maintained under specific pathogen-free conditions following procedures approved by the University of Michigan Committee on the Use and Care of Animals, which are based on the University of Michigan Laboratory Animal Medicine guidelines.

#### *Statistical analyses*

Statistical analyses were performed using GraphPad Prism 9. Two-tailed tests were used for all t-tests. Errors on measurements are represented as standard deviation or standard error of the mean as noted.

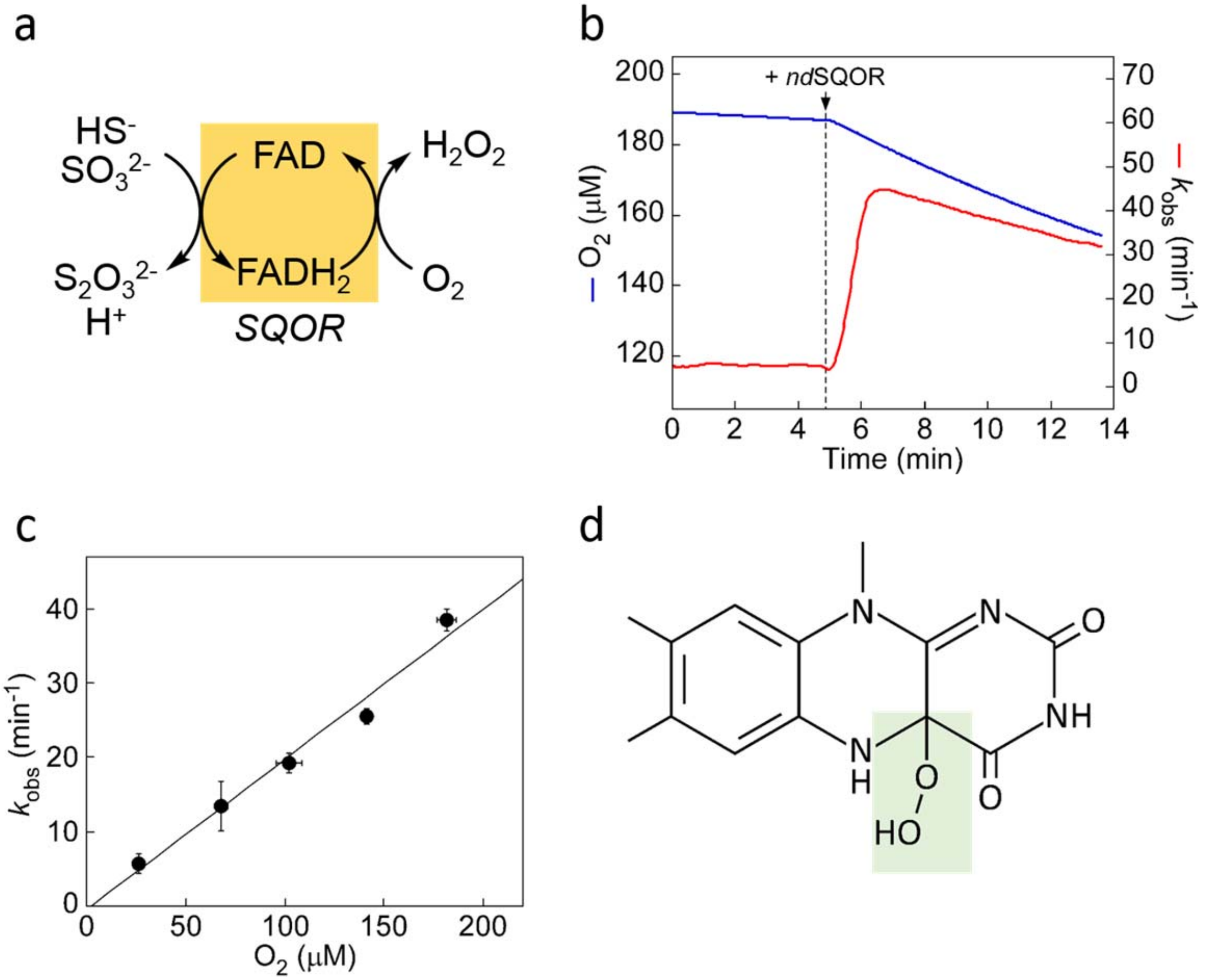


**Supplementary Figure 1. Kinetics of $CoQH_2$ oxidation and $O_2$ consumption catalyzed by *nd*SQOR. (a)** Scheme showing SQOR-catalyzed $H_2O_2$ production. **(b)** Oxygen consumption kinetics in the presence of *nd*SQOR (100 nM) added to a reaction mixture containing sulfide (100 μM) and sulfite (200 μM) in 100 mM potassium phosphate buffer, pH 7.4. **(c)** Dependence of the *nd*SQOR-catalyzed $O_2$ consumption rate on $O_2$ concentration. (**d**) Structure of the proposed 4a-hydroperoxy FAD intermediate. The data are representative (b) or the mean ± S.D. (c) of three independent experiments.

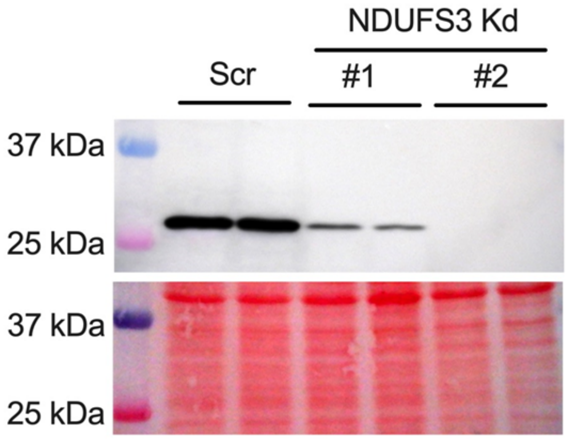


**Supplementary Figure 2. Validation of NDUFS3 knockdown.** Western blot validation of NDUFS3 knockdown with two targeting sequences (#1 and 2, each loaded in duplicate) in HT29 cells. The loading controls (*lower* panel) represents total protein detected with Ponceau S stain.

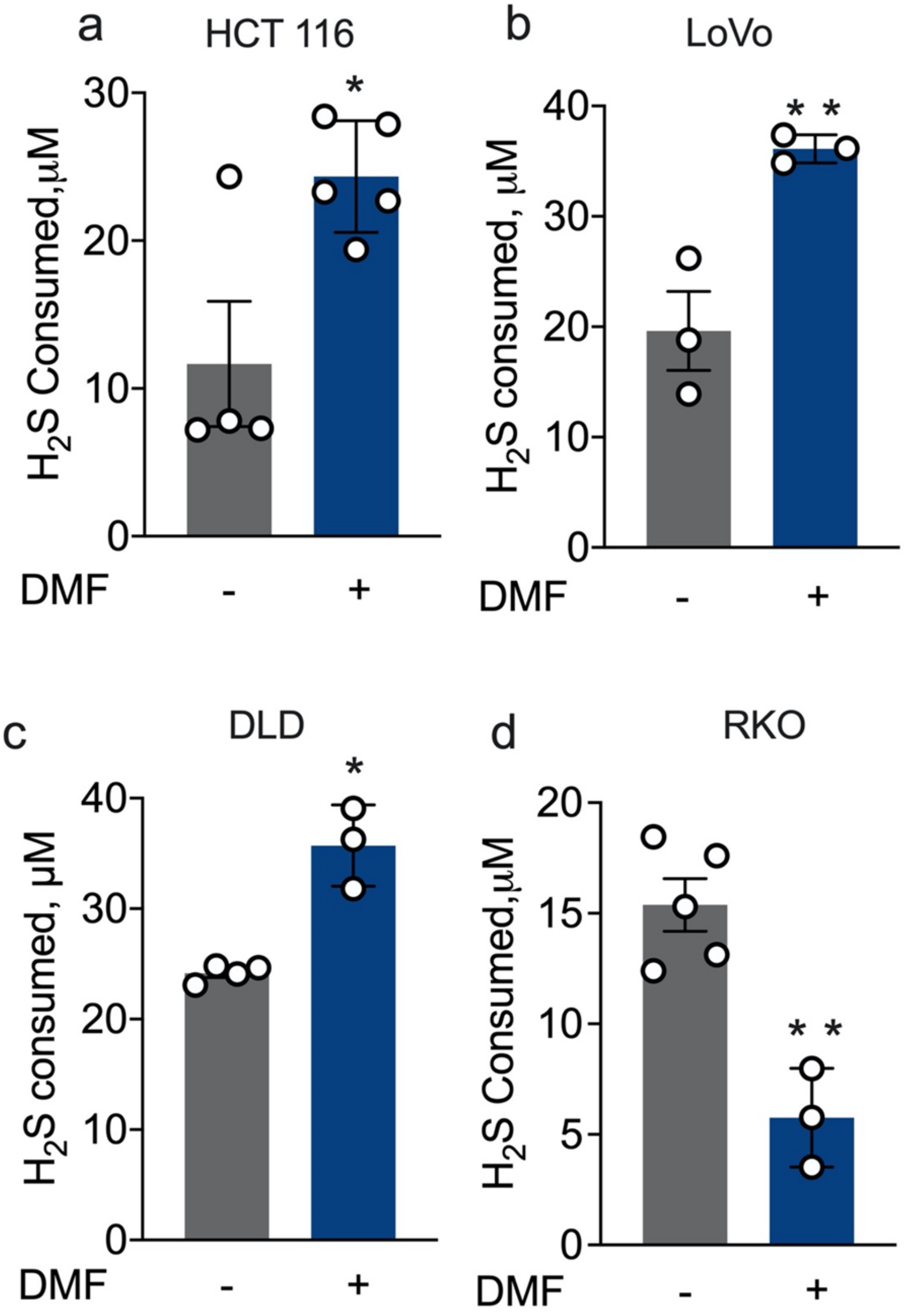


**Supplementary Figure 3. Fumarate acts as terminal electron acceptor in colon cancer cells.** DMF (100 μM) increased $H_2S$ oxidation in **(a)** HCT116, **(b)** LoVo, and **(c)** DLD cells but not in (**d**) RKO cells. The data are the mean ± SEM of 3-5 independent experiments (**$p<0.001$ and *$p<0.05$).

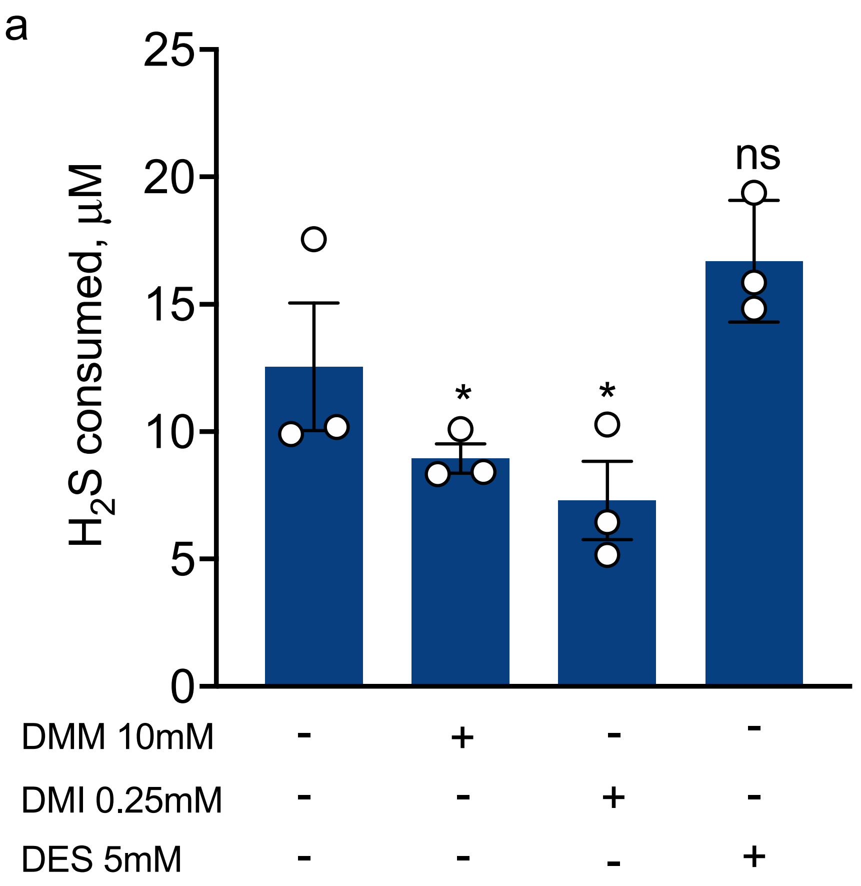


**Supplementary Figure 4. Complex II activity influences $H_2S$ clearance by HT29 cells.** Complex II inhibitors dimethyl malonate (DMM, 10 mM) and dimethyl itaconate (DMI, 0.25 mM) decreased $H_2S$ consumption, whereas diethyl succinate (DES 5 mM) did not show a significant effect. The data represent the SEM of 3 independent experiments (*p<0.01). The control data shown here are the same as for the experiment in Fig. 2c, which was performed in parallel.

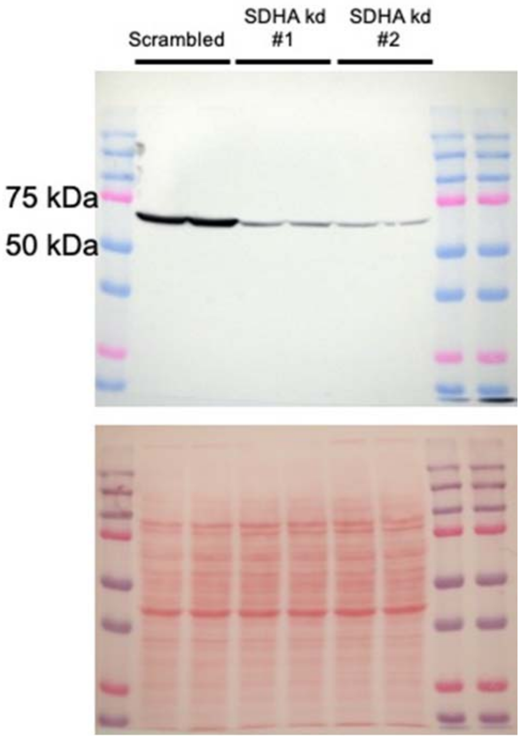


**Supplementary Figure 5. Validation of SDHA knockdown in HT29 cells using two shRNA targeting sequences.** Western blot analysis (*top*) and Ponceau S staining (*bottom*) show expression SDHA in HT29 cells transfected with scrambled or two SDHA targeting sequences (#1 and 2).

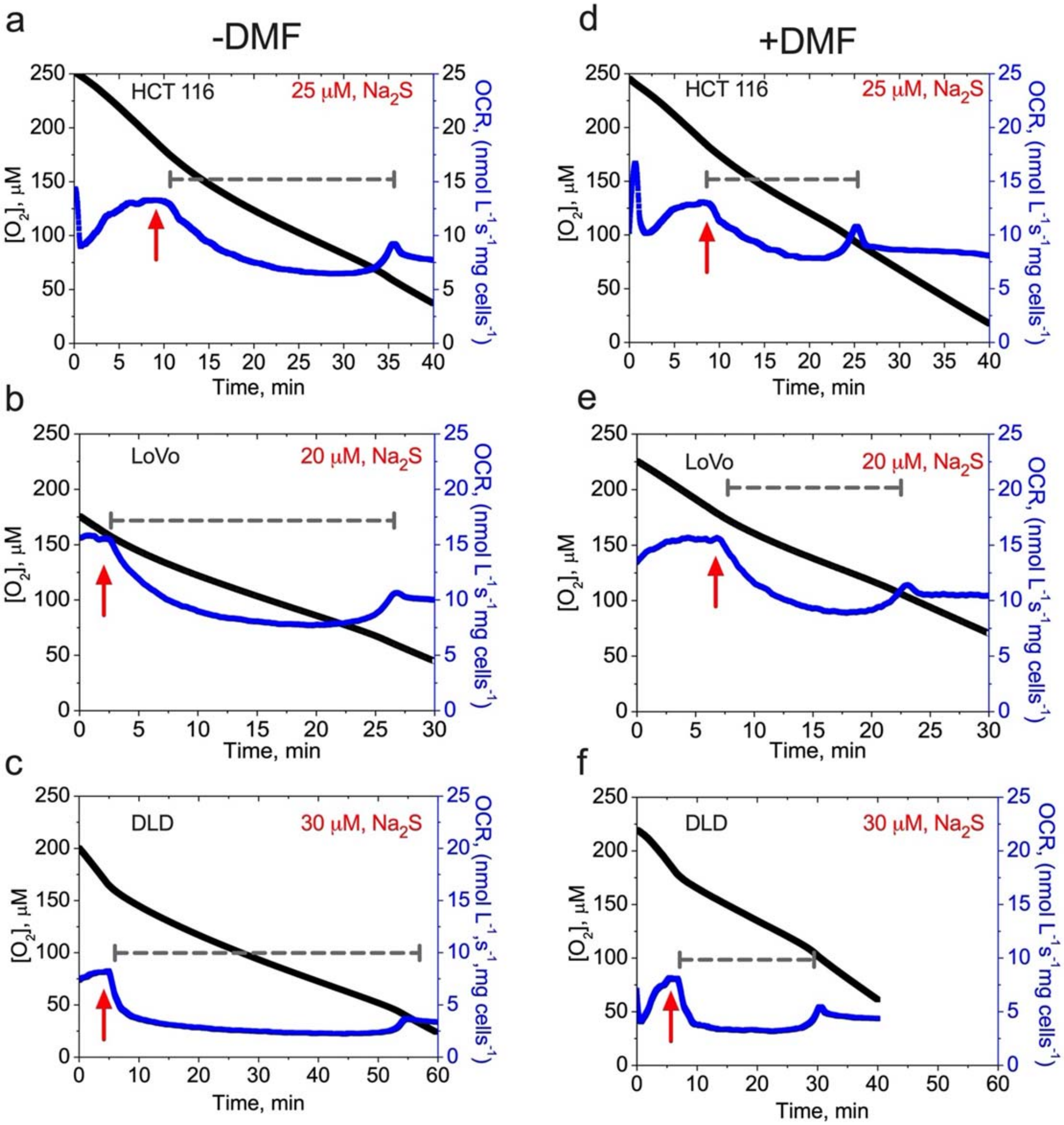


**Supplementary Figure 6. DMF shortens recovery to the basal respiration rate following inhibition by $H_2S$.** Comparison of recovery time to basal OCR following $Na_2S$ exposure in the absence (**a,b,c**) and presence (**d,e,f**) of DMF (200 μM) in three colorectal cancer cell lines (HCT116, LoVo and DLD). The red arrows indicate when $Na_2S$ was added. The length of the gray dashed lines represent the recovery time. The data are representative of 3 independent experiments.

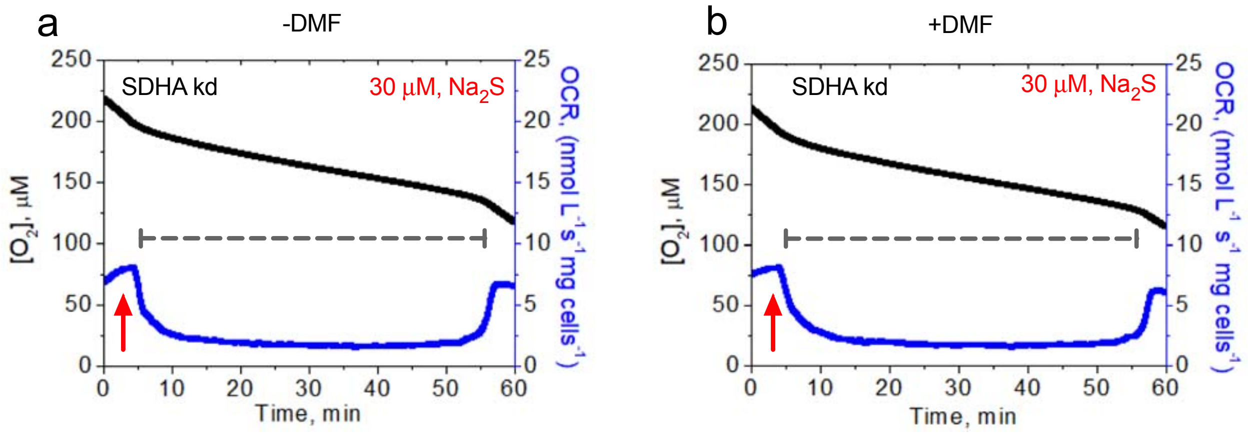


**Supplementary Figure 7. DMF does not affect recovery time to basal respiration rate in SDHA knockdown cells treated with $H_2S$.** Comparison of the time to recovery of basal OCR following $H_2S$ (30 μM) treatment in the absence (**a**) and presence (**b**) of DMF (200 μM) in SDHA knockdown HT29 cells. The red arrows indicate when $Na_2S$ was added. The length of gray dashed lines represents the recovery time. The data are representative of 3 independent experiments.

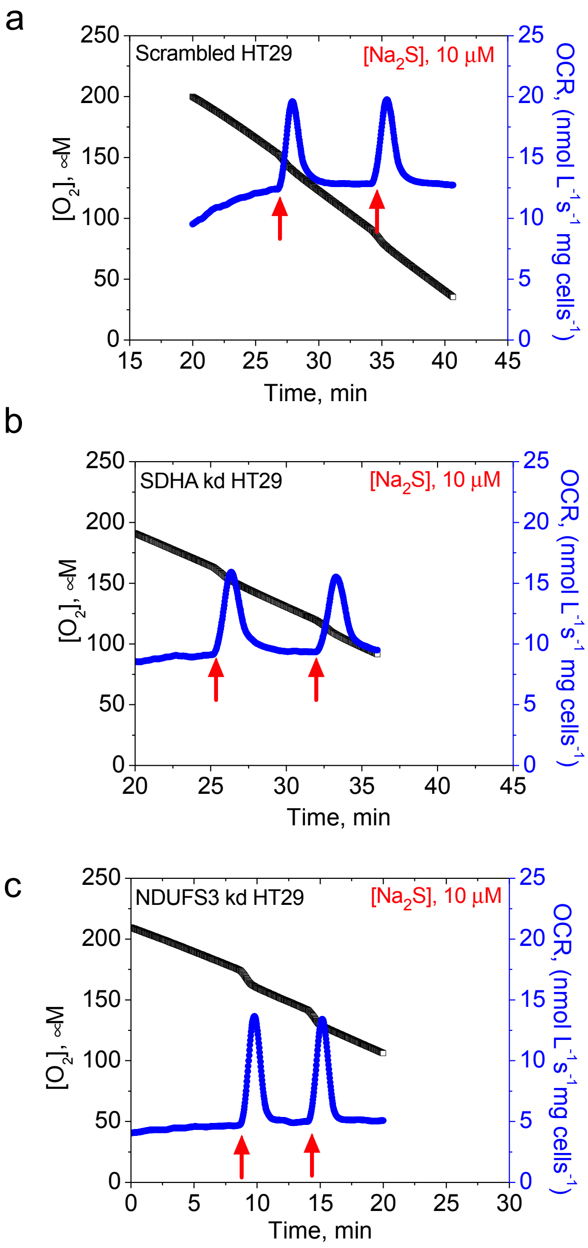


**Supplementary Figure 8. Complex I and II influence $H_2S$-responsive OCR.** Comparison of OCR activation by $H_2S$ (10 μM) in scrambled **(a)**, SDHA knockdown **(b)**, and NDUFS3 knockdown **(c)** HT29 cells. The traces are representative of 3-5 independent experiments. The red arrows indicate when $Na_2S$ was added.

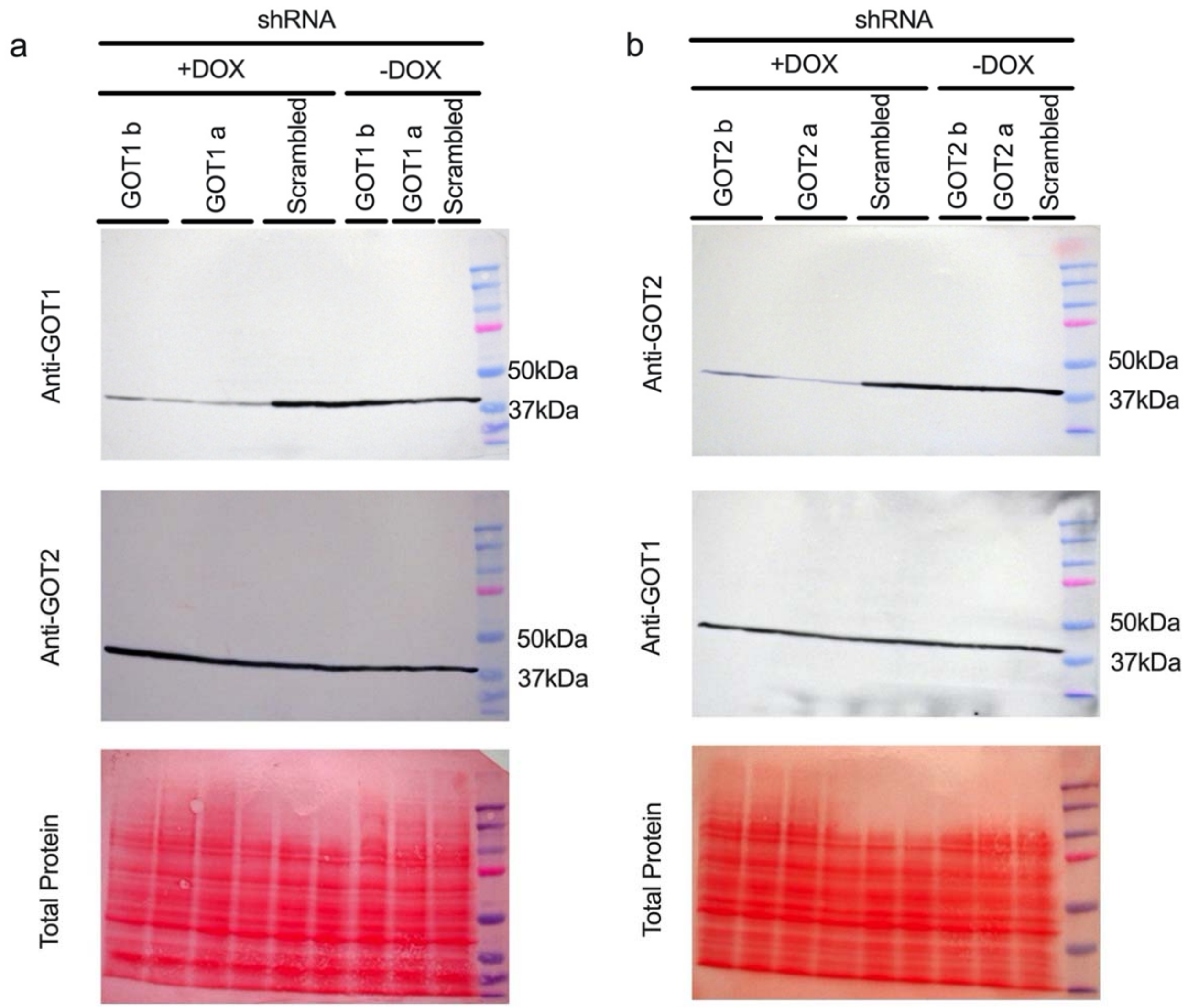


**Supplementary Figure 9. Validation of GOT1 and GOT2 knockdown in HT29 cells. (a)** Expression of GOT1 (*upper*) and GOT2 (*middle*) in GOT1 knockdown cells. **(b)** Expression of GOT2 (*upper*) and GOT1 (*middle*) in GOT2 knockdown cells. Equal loading as indicated by Ponceau S red staining, is shown in the *lower* row in both panels.

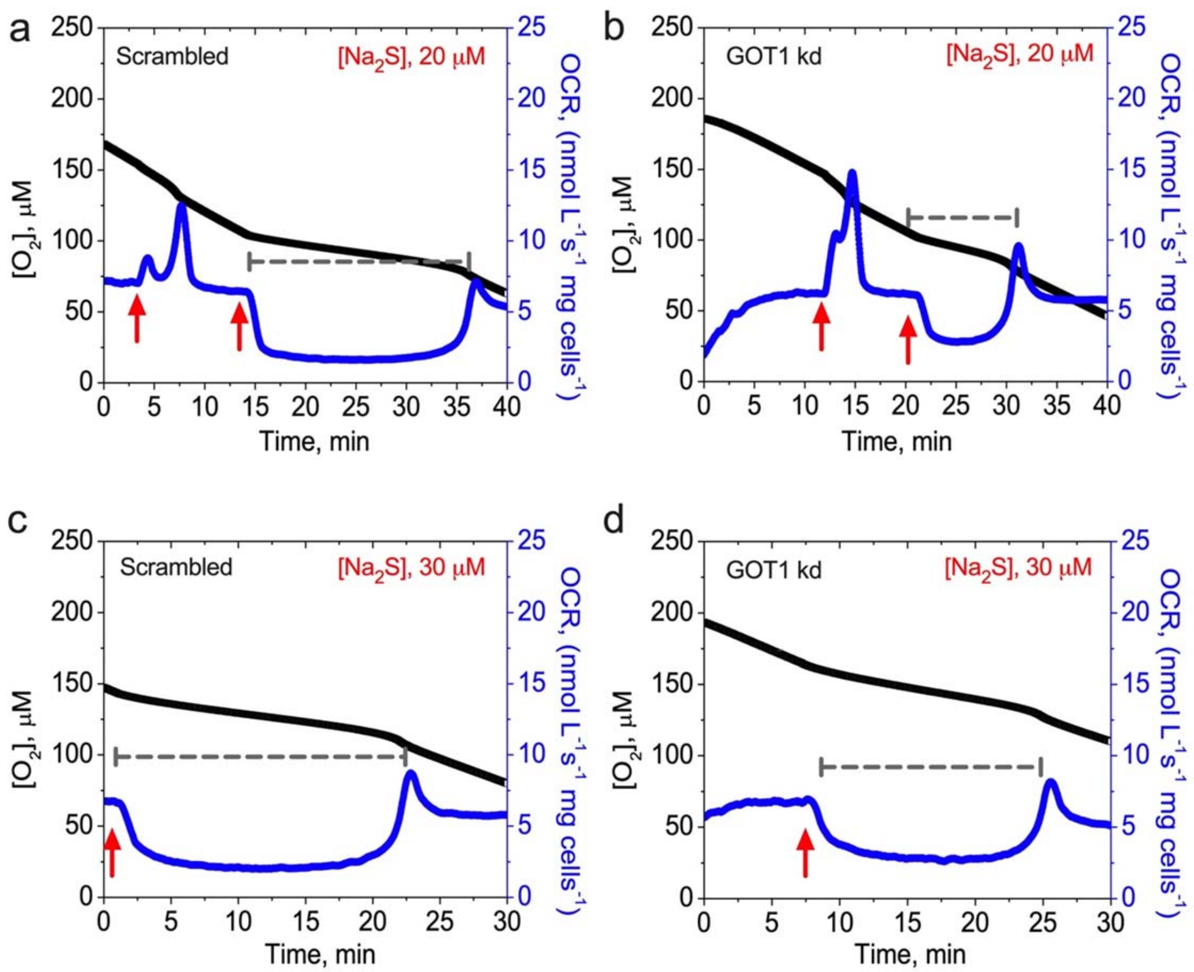


**Supplementary Figure 10. GOT1 knockdown promotes $H_2S$ clearance.** Comparison of the time to recovery of basal OCR following a second injection of $Na_2S$ (20 μM) to (**a**) scrambled versus (**b**) GOT1 knockdown HT29 cells. A similar comparison following injection of a higher concentration of $Na_2S$ (30 μM) to (**c**) scrambled versus (**d**) GOT1 knockdown cells. The red arrows indicate when $Na_2S$ was added. The length of the gray dashed lines represents the recovery time. The data are representative of 3 independent experiments.
